# Preventing large deletions and chromosome loss in engineered human primary T cells by CasPlus with optimized guide RNAs

**DOI:** 10.1101/2025.01.29.635575

**Authors:** Qiaoyan Yang, Jonathan S. Abebe, Michelle Mai, Colin Konishi, Orrin Devinsky, Chengzu Long

## Abstract

Genetically engineered T-cell therapies rely heavily on genome editing tools, such as the CRISPR/Cas9 system. However, unintended on-target chromosomal alterations, including large deletions and chromosome loss can occur and pose significant risks including tumorigenesis. Here we combined CasPlus and optimized guide RNAs to reduce these issues in CRISPR/Cas9 engineering human primary T cells. CasPlus, which integrates an engineered T4 DNA polymerase with Cas9 nuclease and guide RNA, promotes favorable small insertions (1-2 bp) while reducing large deletions and chromosome loss in T cells. Our optimized guide RNAs favoring small insertions reduced large deletions and chromosome loss by two- to five-fold versus those favoring small deletions. Moreover, combining optimized guide RNA with T4 DNA polymerase further synergistically reduced large deletions and chromosome loss by additional two-fold. Notably, replacing currently used guide RNA pairs in clinically applications with optimized pairs biased towards small insertions, along with CasPlus instead of Cas9, for editing greatly reduced large deletions and chromosome loss in gene-edited human primary T cells. These findings demonstrated that pre-selecting target sites favoring small insertions via guide RNA optimization coupled with CasPlus editing is a safer and more effective strategy to improve genome stability in T-cell engineering and other gene-editing applications.

## Introduction

CRISPR/Cas9-mediated genome editing is a powerful tool for gene manipulation, widely utilized in basic research and clinical studies (Cong et al., 2013; Doudna, 2020; Jinek et al., 2012; Li et al., 2023; Mali et al., 2013; Wang and Doudna, 2023). Genetically modified T-cells such as chimeric antigen receptor (CAR) T-cells and T-cell receptor (TCR) engineered T-cells can treat cancers and autoimmune diseases. However, challenges remain regarding safety, efficacy, and immunogenicity (Brentjens et al., 2013; Fesnak et al., 2016; Li et al., 2019; Mackensen et al., 2022; Park et al., 2018; Posey et al., 2024). CRISPR/Cas9-mediated T-cell editing facilitates precise changes in T-cell genomes, increasing CAR T-cell persistence, overcoming CAR T-cell exhaustion, and enabling the generation of therapeutically T cells from healthy donors (Dimitri et al., 2022; Foy et al., 2023; Lu et al., 2020; Stadtmauer et al., 2020; Zhang et al., 2022).

Although initial concerns about CRISPR/Cas9-mediated off-target genome editing have been extensively investigated and mitigated (Bae et al., 2014; Chen et al., 2017; Kleinstiver et al., 2016; Tsai et al., 2017; Wienert et al., 2019; Zhu et al., 2024), unintended chromosomal abnormalities including chromosomal translocations (Stadtmauer et al., 2020), large deletions, and chromosome loss (Leibowitz et al., 2021; Nahmad et al., 2022; Tsuchida et al., 2023) remain insufficiently addressed. These abnormalities could cause complications, including cancer (Beroukhim et al., 2010; Mitelman et al., 2007). This gap persists in preclinical and clinical studies using CRISPR/Cas9 editing of human T-cells. Thus, innovative methods are needed to mitigate chromosomal abnormalities to advance T-cell therapies.

Previously, we developed CasPlus gene editing, comprised of engineered T4 DNA polymerase, Cas9, and guide RNA (gRNA) (Yang et al., 2024). CasPlus enhances small insertions (1–2 bp) and reduces deletions in CRISPR/Cas9-edited HEK293T and induced pluripotent stem cells (iPSCs). In multiplexed gRNAs mediating gene editing for primary human T-cells (All T-cells referred to below are primary unless otherwise specified), CasPlus reduced or eliminated chromosomal translocations. Here we further evaluated the translational potential of CasPlus system to minimize on-target damage in T-cell engineering. Firstly, we confirmed that CasPlus increased small insertions (1–2 bp) while decreasing large deletions in edited human T-cells. By investigating the frequencies of small insertions and large deletions induced by distinct gRNAs, we found that gRNAs favoring small insertions resulted in fewer large deletions versus those favoring small deletions. gRNAs favoring small insertions also significantly decreased chromosome loss, a common problem that can cause aneuploidy and other pathogenic genome instability in edited T-cells (Leibowitz et al., 2021; Nahmad et al., 2022; Tsuchida et al., 2023). Additionally, combining CasPlus editing with gRNA favoring small insertions further minimized the generation of large deletions and chromosome loss. In T-cells edited at multiple loci, CasPlus with optimized gRNA pairs produced small insertions and reduced large deletions and chromosome loss by up to tenfold versus Cas9 with clinical gRNA pairs.

## Results

### CasPlus mitigates on-target large deletions in human T-cells

We firstly conducted a detailed analysis of its impact on small insertions (1-2 bp) and large deletions in human T cells (**Fig. 1A**). We focused on the *TRAC*, *PDCD1*, *CD52* and *B2M* genes, common targets in clinical trials (NCT03399448, NCT04557436) and preclinical research. We utilized the same gRNAs as in these studies for direct comparisons (Liu et al., 2017; Ottaviano et al., 2022; Tsuchida et al., 2023). Activated human T-cells were nucleofected with T4 DNA polymerase mRNA/Cas9 mRNA/gRNAs (CasPlus editing) or Cas9 mRNA/gRNAs (Cas9 editing) complexes. Four days post-nucleofection, a pool of edited cells was collected. The targeted region of ∼300 base pair (bp) or 5 kilobases (kb) was amplified by PCR for analysis. Amplicons of ∼300 bp were analyzed by deep sequencing to evaluate the alteration of small insertions (1-2 bp) and small deletions (1–100 bp). Amplicons of ∼5 kb were investigated by long-read PacBio sequencing, with reads smaller than 4.5 kb (< 4.5 kb) were classified as large deletions (500 bp < Deletion size < 5 kb) (**Fig. 1B**). CasPlus significantly enhanced 1 bp insertions and reduced >1 bp deletions across four of six gRNAs tested in T-cells (**Fig. 1C**). To exclude non-biological PCR and PacBio sequencing artifacts, we normalized PacBio read depth by dividing the number of PacBio sequencing read depth at each nucleotide position by the total number of PacBio reads (**Supplemental Fig. S1A**). PacBio sequencing showed an increase in normalized depth coverage across the entire PCR amplicons in CasPlus-edited verse Cas9-edited cells across all four gRNAs (**Fig. 1D**). CasPlus increased the frequency of reads from 4.5 kb to 5 kb, while reducing those < 4.5 kb (**Fig. 1E** and **supplemental Fig. S1B and table S1).** With two additional gRNAs, PDCD1-g1 and B2M-g5, CasPlus increased 1 bp deletions while decreasing >1 bp deletions (**Supplemental Fig. S1C**). The frequency of reads in the 4.5 kb to 5 kb range was comparable between CasPlus-versed Cas9-edited T-cells for these gRNAs (**Supplemental Fig. S1D-1E and table S1**), suggesting that CasPlus is less efficient in preventing large deletions with gRNAs favoring small deletions versus Cas9. CasPlus with gRNA favoring small insertions achieved an ∼2-fold reduction in large deletions versus Cas9 editing in T-cells (**Fig. 1F**). The engineered RB69 DNA polymerase, derived from a T4-related bacteriophage, functions similarly with T4 DNA polymerase in HEK293T cells (Yang et al., 2024). RB69 DNA polymerase-mediated CasPlus consistently produced similar alterations in small indel profile and large deletions with T4 DNA polymerase-mediated CasPlus in T cells (**Supplemental Fig. S1F-1H and table S1**).

**Fig. 1.**
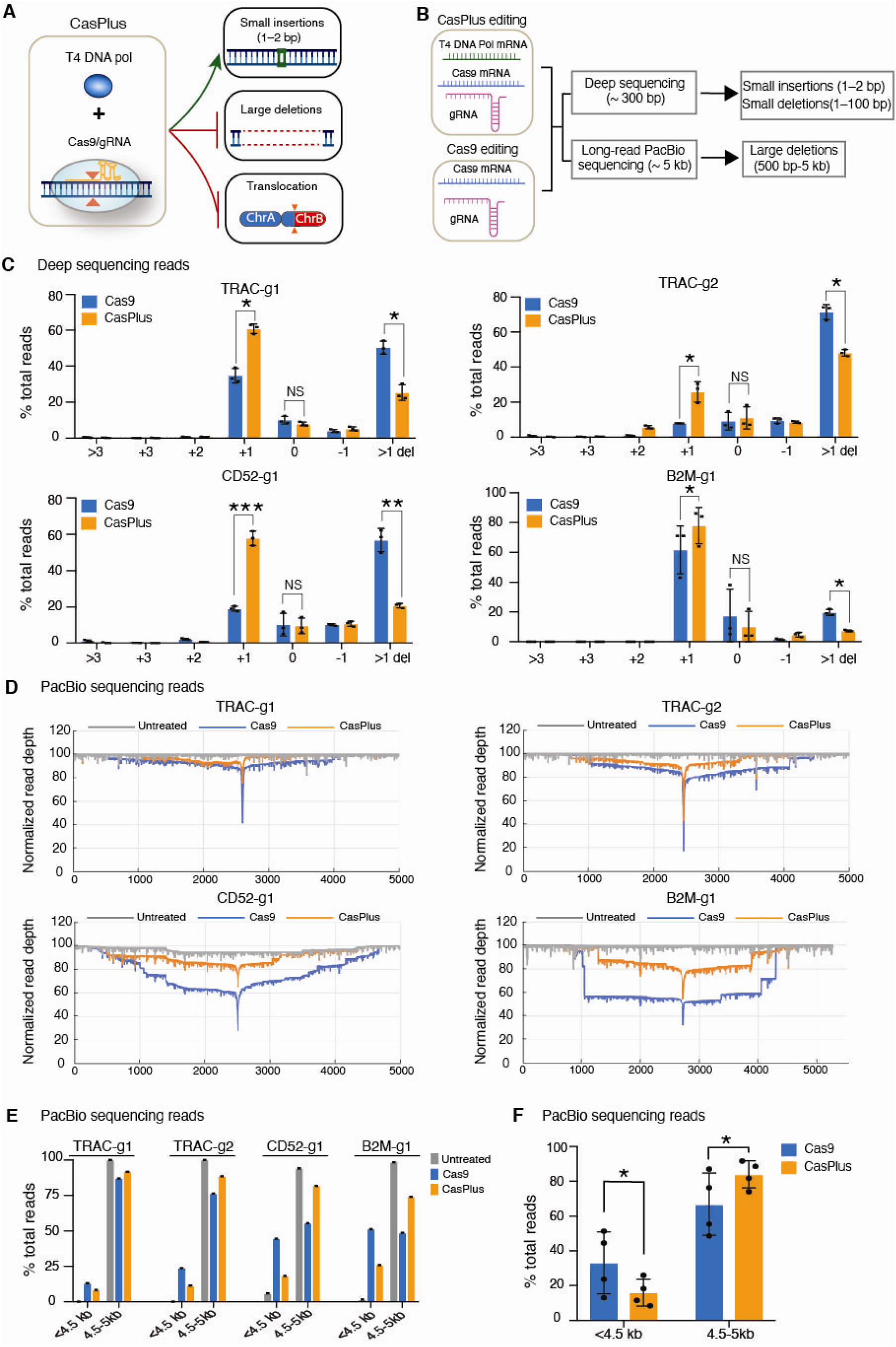
CasPlus mitigates on-target large deletions in human primary T cells. **(A)** Previous study (Yang et al., 2024) demonstrated that CasPlus enhances the frequency of small insertions (1–2 bp), decreases the occurrence of small and large deletions, and prevents chromosomal translocations in CRISPR/Cas9-edited HEK293T and iPS cells. (**B**) A schematic illustrating the workflow for analyzing small indels and large deletions in T cells. T cells were nucleofected with T4 DNA polymerase mRNA/Cas9 mRNA/gRNA or Cas9 mRNA/gRNA and collected four days post-nucleofection. Targeted region (300 bp or ∼5 kb) was amplified and investigated by short-read deep sequencing to assess small indels or long-read PacBio sequencing to discover large deletions. PacBio read depth was analyzed based on the pipeline described in the **Methods**. Reads with sizes < 4.5 kb were considered as large deletions. (**C**) Frequencies of indels at *TRAC*, *CD52* or *B2M* loci in T cells nucleofected with TRAC-g1, TRAC-g2, CD52-g1 or B2M-g1 for Cas9 and CasPlus editing. Data was shown as mean ± SEM. Each dot represents one biological replicate. *, p<0.05; **, p<0.01; ***, p<0.005; ****, p<0.001. (**D**) Normalized depth of PacBio reads at *TRAC*, *CD52* or *B2M* loci in untreated, Cas9-edited, and CasPlus-edited T cells with TRAC-g1, TRAC-g2, CD52-g1 or B2M-g1, respectively. (**E**) Quantification of PacBio reads with sizes less than 4.5 kb or with sizes of 4.5 kb to 5 kb in samples described in (**D**). Each dot represents one biological replicate. (**F**) Overall frequency of PacBio reads with size less than 4.5 kb or with size of 4.5 kb to 5 kb across four gRNAs in Cas9- and CasPlus-edited T cells. Data was shown as mean ± SEM.

### Combing gRNA favoring small insertions and CasPlus minimizes on-target large deletions

Cas9-induced on-target small insertions and large deletions are influenced by the gRNA context (Allen et al., 2018; Kosicki et al., 2022; Leenay et al., 2019; Owens et al., 2019; Shen et al., 2018), but the relationship between these outcomes in Cas9- and CasPlus-edited T cells remains unclear. To explore this, we designed a pool of gRNAs to target the first coding exons of *TRAC*, *PDCD1*, *CD52* and *B2M* genes (4-6 gRNAs for each gene) and compared their efficiency in inducing small insertions and large deletions. Consistent with prior reports (Allen et al., 2018; Leenay et al., 2019; Shen et al., 2018; Yang et al., 2024), the frequencies of small insertions induced by Cas9 or CasPlus systems varied depending on the gRNA context (**Supplemental Fig. S2A**). Among six gRNAs targeting gene *TRAC*, TRAC-g3 was the most efficient at inducing small insertions (+1bp) whereas TRAC-g5 produced minimal small insertions (**Fig. 2A**). In Cas9-treated T-cells, TRAC-g3 reduced PacBio reads with large deletions 5-fold versus TRAC-g5 [TRAC-g3 (7.6%); TRAC-g5 (37.9%)] (**Fig. 2B-2C and supplemental Fig. S2B**). CasPlus with TRAC-g3 reduced large deletions to additional 2-fold compared to Cas9 with TRAC-g3 [(CasPlus (3.8%); Cas9 (7.6%)] (**Fig. 2B-2C and supplemental Fig. S2B**). For five gRNAs targeting the PDCD1 gene, PDCD1-g2 was most efficient while PDCD1-g1 was the least efficient in producing small insertions (+1bp) (**Fig. 2D**). Cas9 with PDCD1-g2 showed a 2-fold reduction in large deletions versus PDCD1-g1 [PDCD1-g2 (24.4%); PDCD1-g1 (46.5%)] (**Fig. 2E-3F and supplemental Fig. S2C**). CasPlus with PDCD1-g2 reduced large deletions by an additional 2-fold versus Cas9 [(CasPlus (10.8%); Cas9 (24.4%)] (**Fig. 2E-3F and supplemental Fig. S2C**). Thus, gRNAs favoring small insertions resulted in 2-to-5-fold fewer large deletions. Further, pairing these optimized gRNAs with CasPlus reduces large deletions an additional 2-fold.

**Fig. 2.**
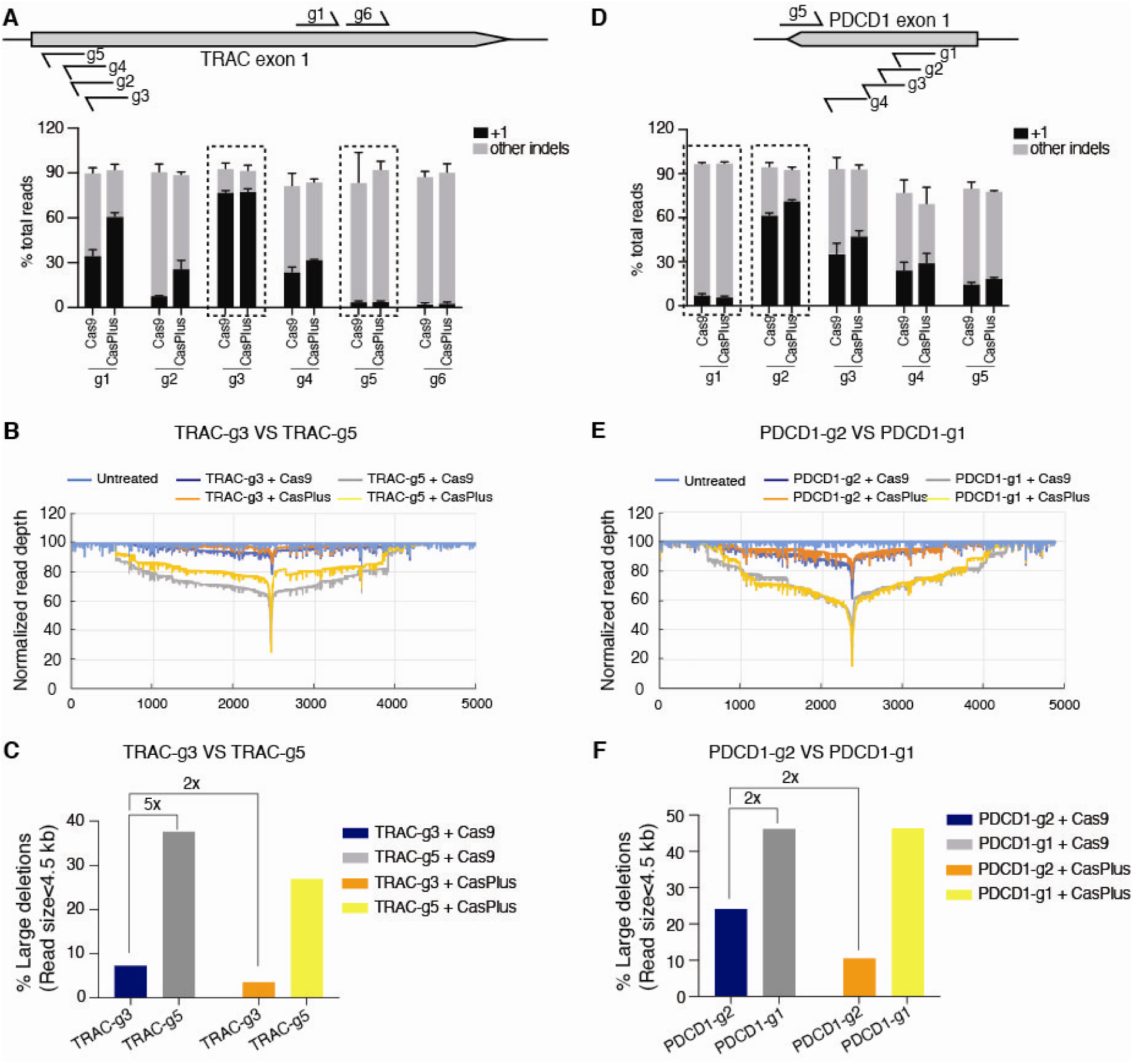
Combining gRNAs favoring small insertions and CasPlus minimizes on-target large deletions. **(A)** Frequencies of 1-bp insertions or other indels observed in T cells with one single gRNA targeting TRAC exon 1 for Cas9 or CasPlus editing. Data was shown as mean ± SEM from three biological replicates. (**B**) Normalized depth of PacBio reads at *TRAC* locus in untreated T cells or T cells with TRAC-g3 or TRAC-g5 for Cas9 and CasPlus editing. (**C**) Quantification of PacBio reads with large deletions (size < 4.5 kb) in samples described in panel (**B**). **(D)** Frequencies of 1-bp insertions or other indels observed in T cells with one single gRNA targeting PDCD1 exon 1 for Cas9 or CasPlus editing. Data was shown as mean ± SEM from three biological replicates. (**E**) Normalized depth of PacBio reads at *PDCD1* locus in untreated T cells or T cells with PDCD1-g2 or PDCD-g1 for Cas9 and CasPlus editing. (**F**) Quantification of PacBio reads with large deletions (size < 4.5 kb) in samples described in panel (**E**).

**Fig. 3.**
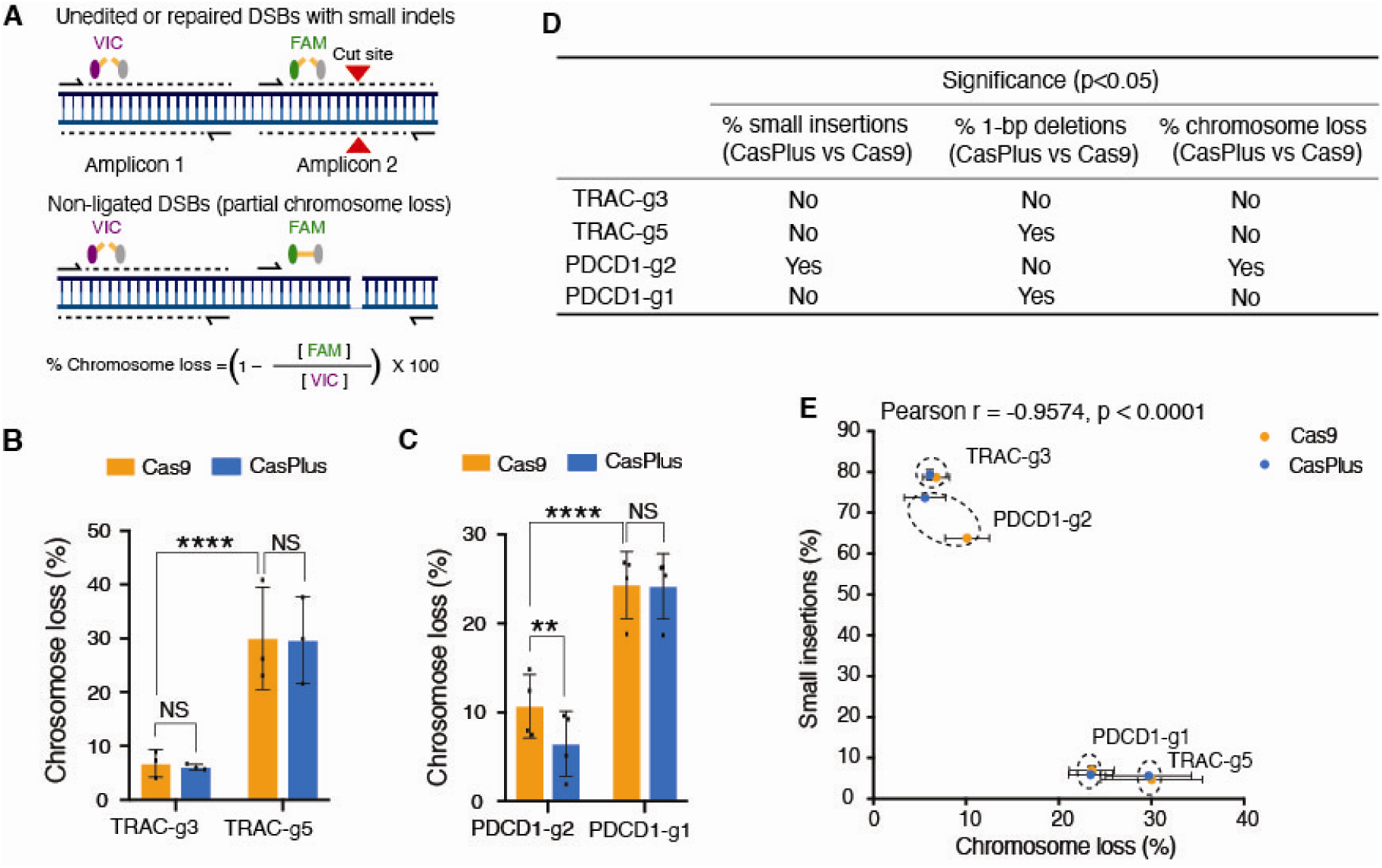
Combining gRNAs favoring small insertions and CasPlus minimizes chromosome loss. **(A)** Schematic of ddPCR assay to measure partial chromosome loss. The red triangles represent the predicted cut site. The detection of both VIC and FAM probes indicates no DSBs or repaired DSBs with small indels (**Upper panel**). The detection of VIC but not the FAM probe indicated non-ligated DSBs (**Bottom panel**). **(B)** Quantifications of partial chromosome loss measured by ddPCR at *TRAC* locus in T cells with TRAC-g3 or TRAC-g5 for Cas9 and CasPlus editing. Data was shown as mean ± SEM. Each dot represents one biological replicate. **(C)** Quantifications of partial chromosome loss measured by ddPCR at *PDCD1* locus in T cells with PDCD1-g2 or PDCD1-g1 for Cas9 and CasPlus editing. Data was shown as mean ± SEM. Each dot represents one biological replicate. (**D**). Table showing the presence (Yes) or absence (No) of significant alterations (p<0.05) in frequency of small insertion, 1-bp deletions, or chromosome loss between Cas9- and CasPlus-edited T cells with one single gRNA targeting gene *TRAC* or *PDCD1*. (**E**) Correlation between the rate of chromosome loss measured by ddPCR and the frequency of small insertions measured by deep sequencing.

### Combining gRNAs favoring small insertions and CasPlus minimizes chromosome loss

To assess the impact of gRNA sequences and CasPlus on chromosome loss, we used the validated multiplexed droplet digital PCR (ddPCR) assay (Tsuchida et al., 2023). Two ∼200 bp amplicons were designed for each target gene (**Fig. 3A**). Amplicon 1, 200-300 bp away from the predicted Cas9 cleavage site, included a probe labelled with fluorescent dye VIC. Amplicon 2, labelled with 6-fluorescien (FAM), spanned the Cas9 cleavage site, anticipating a DSB without re-ligation would inhibit amplicon 2 amplification. The primers and probes for amplicon 2 were positioned to avoid binding-site disruption by small insertions or deletions, the most common and intended edits after Cas9 editing (**Supplemental Fig. S3A-3B**). Detection of amplicon 1 (control amplicon) without detection of amplicon 2 (targeted amplicon) was interpreted as partial chromosome loss (Tsuchida et al., 2023).

In edited T-cells, Cas9 resulted in 4.4-fold reduction in chromosome loss with TRAC-g3 (favoring +1 bp) versus TRAC-g5 (favoring >1 bp deletions) [TRAC-g3 (6.8%); TRAC-g5 (30.0%)]; CasPlus did not significantly alter chromosome loss versus Cas9 with TRAC-g3 [CasPlus (6.1%); Cas9 (6.8%)] (**Fig. 3B**). For PDCD1, Cas9 reduced chromosome loss 2.3-fold with PDCD1-g2 (favoring +1 bp) versed PDCD1-g1 (favoring >1 bp deletions) [PDCD1-g2 (10.7%); PDCD1-g1 (24.3%)]; CasPlus with PDCD1-g2 decreased chromosome loss by additional 1.6-fold compared to Cas9 [(CasPlus (6.5%); Cas9 (10.7%)] (**Fig. 3C**). CasPlus significantly increased the frequency of 1-bp deletions over >1 bp deletions for TRAC-g5 or PDCD1-g1 (**Supplemental Fig. S3C-3D**) but did not alter chromosome loss rate induced by these gRNAs (**Fig. 3B-3C**). CasPlus mitigation of chromosome loss strongly correlated with an increase in small insertions over small deletions (**Fig. 3D**). This correlation was confirmed by Pearson correlation coefficient analysis, with a negative correlation between the rates of chromosome loss and frequencies of small insertions (Pearson correlation = −0.9547, p<0.001) (**Fig. 3E**). These results demonstrate that chromosome loss can be reduced through gRNA optimization and further minimized by combination with CasPlus editing.

### Combining optimized gRNA pairs with CasPlus mitigated large deletions and chromosome loss in multiplexed gene-edited T cells

Since gRNAs favoring small insertions reduce the likelihood of large deletions and chromosome loss, we hypothesized that replacing clinical gRNAs with optimized gRNAs greatly favoring small insertions would produce T-cells with fewer chromosomal abnormalities. To test this hypothesis, we nucleofected T cells with clinical gRNA combinations of TRAC-g1 and PDCD1-g1 or the optimized gRNA combinations of TRAC-g3 and PDCD1-g2 for Cas9 and CasPlus editing. The optimized gRNA combination of TRAC-g3 and PDCD1-g2 induced higher frequencies of small insertions without compromising overall editing efficiency at TRAC or PDCD1 individual sites compared to clinical gRNA combination (**Fig. 4A and supplemental Fig. S4A-4B**). In PacBio reads for Cas9 treated T-cells, we observed a 4.9-fold decrease in large deletions at TRAC site [(TRAC-g3 (9.2%); TRAC-g1 (44.9%)] and 4.1-fold reduction at PDCD1 site [(PDCD1-g2 (16.6%); PDCD1-g1 (67.5%)] with gRNAs TRAC-g3/PDCD1-g2 compared to TRAC-g1/ PDCD1-g1 (**Fig 4B-4C** and **supplemental Fig. S4C)**). CasPlus with optimized gRNA combinations reduced the large deletions to additional 2.2-fold at TRAC site [CasPlus (4.3%); Cas9 (9.2%)] and 1.3-fold at PDCD1 site [(CasPlus (13.0%); Cas9 (16.6%)] versus Cas9 (**Fig 4B-4C** and **supplemental Fig. S4C**).

**Fig. 4.**
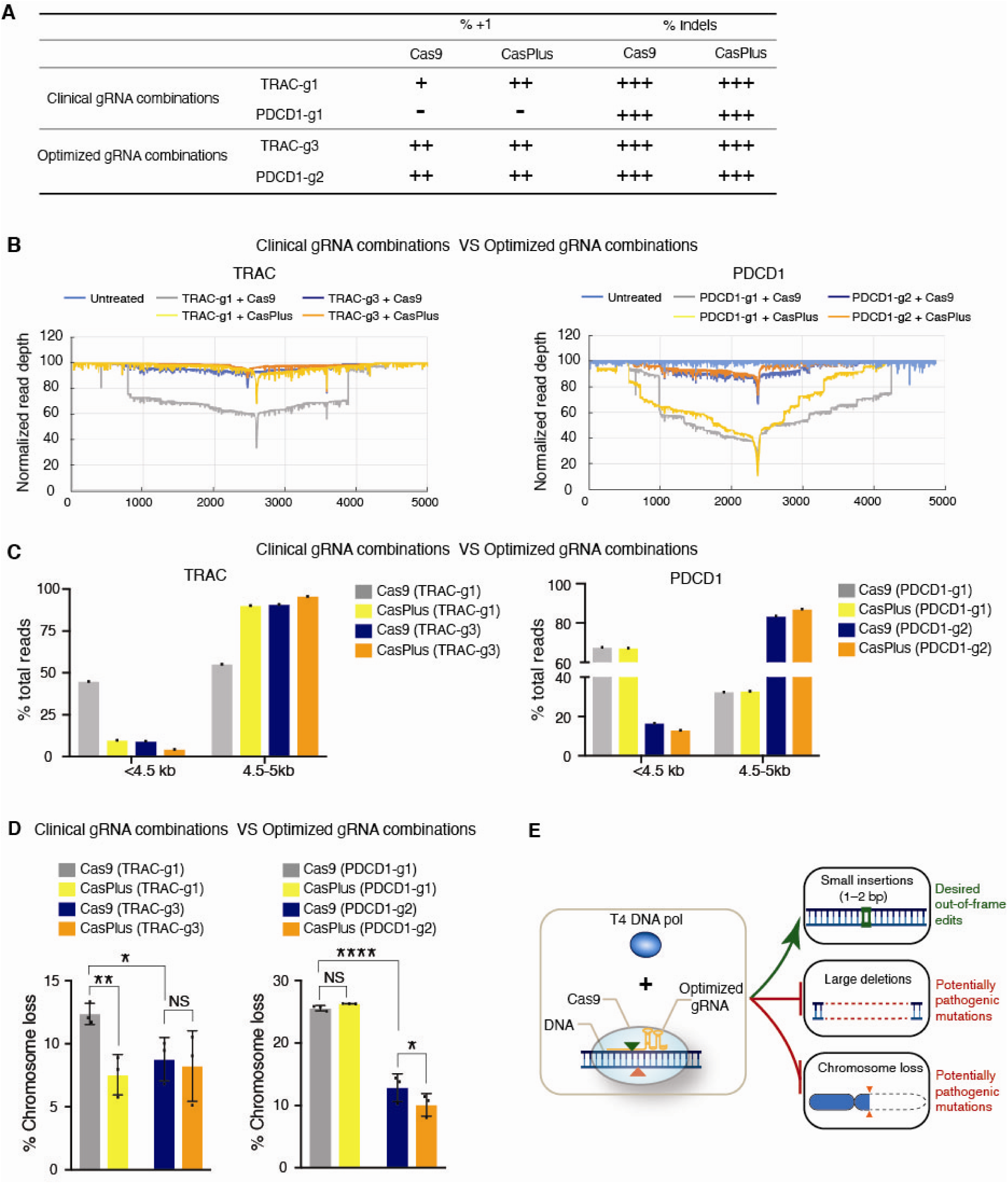
**Combining optimized gRNA pairs with CasPlus mitigated large deletions and chromosome loss in multiplexed gene-edited T cells.** (A). Table showing frequencies of 1-bp insertion and total indels in T cells induced by TRAC-g1 and PDCD1-g1 or TRAC-g3 and PDCD1-g2, alone with mRNA for Cas9 or CasPlus editing. One, two, three plus symbols (+) represent frequencies of ∼30%, ∼60%, and ∼90% respectively. (**B**) Normalized depth of PacBio reads at *TRAC* (**Left panel**) and *PDCD1* (**Right panel**) loci in untreated T cells or multiplexed gene-edited T cells with clinical used gRNA combinations (TRAC-g1 + PDCD1-g1) or optimized gRNA combinations (TRAC-g3 and TRAC-g2) for Cas9 and CasPlus editing. (**C**) Quantification of PacBio reads with sizes less than 4.5 kb or with sizes of 4.5 kb to 5 kb in samples described in (**B**). Each dot represents one biological replicate. (**D)** Quantifications of partial chromosome loss measured by ddPCR at *TRAC* (**Left panel)** and *PDCD1* (**Right panel)** loci in multiplexed gene-edited T cells with TRAC-g3 and PDCD1-g2 or TRAC-g1 and PDCD1-g1 for Cas9 and CasPlus editing. Data was shown as mean ± SEM. Each dot represents one biological replicate. (**E)** Schematic illustrating the features of CasPlus editing. CasPlus editing with gRNA predominantly favoring small insertions can effectively increase small insertions (1–2 bp) and decrease potentially pathogenic outcomes including large deletions and chromosome loss in engineered human primary T cells.

Chromosome losses are also decreased with optimized gRNAs. Cas9 with TRAC-g3 and PDCD1-g2 resulted in a 1.4-fold reduction in chromosome loss at TRAC site [(TRAC-g3 (8.8%), TRAC-g1 (12.4%) and 2.0-fold reduction at PDCD1 site [PDCD1-g2 (12.8%), PDCD1-g1 (26.2%)] compared to Cas9 with clinically used gRNAs (**Fig. 4D**). CasPlus, while not significantly altering chromosome loss at TRAC site with TRAC-g3, reduced chromosome loss an additional 1.3-fold at PDCD1 site with PDCD1-g2 compared to Cas9 [(CasPlus (9.5%), Cas9 (12.8%)] (**Fig. 4D**). Therefore, we propose pre-selecting target site favoring small insertions and pairing it with CasPlus editing to minimize large deletions and chromosome loss in multiplexed gene-edited human T-cells.

## Discussion

Current CRISPR gRNA design algorithms primarily focus on minimizing off-target effects and enhancing on-target editing efficiency by considering sequence features and genome context. However, their accuracy and safety remain limited by an incomplete understanding of repair outcomes, indel biases, variable chromatin accessibility, and cell-type-specific differences. Here, we demonstrate that the CasPlus gene editing system, combined with optimized gRNAs, promotes favorable small insertions (1-2 bp) while significantly reducing potentially pathogenic edits, including large deletions and chromosome loss, induced by conventional CRISPR/Cas9 editing in human primary T cells (**Fig. 4E**). This advancement in genetic engineering offers a safer and more efficient strategy with reduced genomic toxicity for T-cell therapies and other gene-editing applications.

Identifying the frequency of CRISPR/Cas9 induced small indels and large deletions remains challenging using a single method. Short-read deep sequencing can effectively detect small indels but lacks the capability to identify large deletion, while long-read PacBio sequencing effectively detects large deletions but lacks the precision to accurately assess small indels (Hu et al., 2021). Our findings revealed a positive correlation between increased small insertions measured through deep sequencing and reduced large deletions detected by PacBio sequencing in edited T-cells, consistent with recent discoveries (Hwang et al., 2024). Thus, we propose using deep sequencing or Sanger sequencing to pre-select target sites favoring small insertions to minimize the likelihood of large deletions.

Moreover, our analysis of template-free Cas9 editing outcomes highlighted the importance of nucleotide preferences in gRNA design. Specifically, thymine/adenine (T/A) at position 17 (5 nt upstream of the PAM) within the gRNA sequence tends to favor small insertions(Allen et al., 2018; Leenay et al., 2019; Shen et al., 2018). Systematic profiling of Cas9 incision also links the configuration of Cas9 cuts with precise and predicable 1-bp insertions, with guanine (G) at position 18 being highly associated with staggered double-strand DNA breaks (DSBs) and small insertions (Longo et al., 2024). Therefore, integrating these nucleotides preferences at positions 17 and 18 during gRNAs design would facilitate the optimization process for both Cas9 and CasPlus editing.

Digital droplet PCR (ddPCR) and single cell RNA sequencing (scRNA-seq) are established methods for measuring chromosome loss in T-cells (Nahmad et al., 2022; Tsuchida et al., 2023). While ddPCR can only primarily detect partial rather than whole chromosome loss, a previous comprehensive study demonstrated a positive correlation between ddPCR and scRNA-seq assays, supporting ddPCR as reliable tool for this purpose (Tsuchida et al., 2023). In our analysis, ddPCR revealed that CasPlus reduced chromosome loss compared to Cas9 with gRNA favoring small insertions. However, CasPlus was unable to reduce chromosome loss with gRNA predominantly favor small deletions, despite its ability to preferentially enhance 1-bp deletion over >1 deletions at these sites. Importantly, our analysis revealed a strong negative correlation between chromosome loss and the frequency of small insertions, underscoring the potential of gRNA optimization as an effective strategy to decrease chromosome loss.

Alternative approaches to limit chromosome loss have been actively explored. For instance, modified T-cell preparation protocol delivers Cas9 editing components into non-activated T cells to reduce chromosomal abnormalities (Tsuchida et al., 2023). However, non-activated T cells, characterized by condensed chromatin and a resting state, exhibit reduced accessibility for CRISPR/Cas9, leading to lower editing efficiency (Verkuijl and Rots, 2019). Moreover, DNA repair efficiency in non-activated T cells is significantly lower than in activated T cells, leading to a higher accumulation of unrepaired lesions and increased cell death (Hu et al., 2018). Emerging CRISPR/Cas9-based technologies, such as base and prime editing, which do not generate double-strand break (DSBs), hold potential to avoid high levels of chromosomal abnormalities. However, base editing can only induce one or a few nucleotide substitutions and has off-target effect on both DNA and RNA(Gaudelli et al., 2017; Komor et al., 2016). Prime editing allows a broader range of edits(Anzalone et al., 2019), but has lower average editing efficiency compared to Cas9 (Kim et al., 2021), which limits its potential multiplexed gene-editing applications. Despite these advancements, CRISPR/Cas9 remains the primary platform for clinical applications due to its higher editing efficiency and broader range of potential modifications. Our study highlights a simple yet effective method to mitigate unintended chromosomal abnormalities by incorporating T4 DNA polymerase and optimizing gRNA sequences within current engineered T cell manufacturing pipelines.

## Methods

### Plasmids

The vector pSpCas9(BB)-2A-GFP (PX458) (Addgene plasmid #48138) containing the human codon-optimized SpCas9 gene with 2A-GFP and the sgRNA backbone and vector pCMV-T7-SpRY-P2A-EGFP (RTW4830) (Addgene plasmid #139989) were purchased from Addgene. EF1A-MS2-T4-2A-EGFP and EF1A-MS2-RB69-2A-EGFP was constructed as described previously(Yang et al., 2024). The pCMV-T7-SpCas9, pCMV-T7-T4, pCMV-T7-RB69 were generated by amplifying the SpCas9, T4 and RB69 from pSpCas9(BB)-2A-GFP (PX458), EF1A-MS2-T4-2A-EGFP and EF1A-MS2-RB69-2A-EGFP, respectively, and subcloning of these PCR fragments into digested pCMV-T7-SpRY-P2A-EGFP (RTW4830) backbone via Gibson assembly.

### In vitro transcription

The Cas9 mRNA (5meC, Ψ) was purchased from TriLink Biotechnologies (L-6125) and gRNAs against genes *TRAC*, *PDCD1*, *CD52* or *B2M* were synthesized from Synthego (https://ice.synthego.com/<u>#/</u>). The vectors pCMV-T7-SpCas9, pCMV-T7-T4, pCMV-T7-RB69 were used as template for in vitro transcription using Invitrogen™ mMESSAGE mMACHINE™ T7 ULTRA Transcription Kit (ThermoFisher Scientific) according to the manufacturer’s instruction. The raw RNA transcripts were purified using MEGAclear Transcription Clean Up Kit (ThermoFisher Scientific) and eluted with nuclease-free water (Ambion). Concentration of Cas9, T4 or RB69 DNA polymerase mRNA was measured by a NanoDrop instrument (Thermo Scientific). Synthesized gRNAs are listed in **Supplemental table S2**.

### Human T-cell culture, isolation, and stimulation

Human peripheral blood mononuclear cells (PBMCs) were purchased from Lonza. Frozen PBMCs were thawed and cultured in X-vivo 15 medium (Lonza) supplemented with 5% human AB serum (Heat inactivated) (Valley Biomedical) for 24 hours. T cells were isolated from human PBMCs using EasySep Human T Cells Isolation Kits (StemCell Technologies) according to the manufacturer’s instructions. Immediately after isolation, T cells were activated and stimulated with a 1:1 ratio of anti-human CD3/CD28 magnetic Dynabeads (ThermoFisher) to cells in X-vivo 15 medium supplemented with 5% human AB serum (Heat inactivated), 5 ng/ml IL-7 (PeproTech), 5 ng/ml IL-15 (PeproTech), and 200 U/ml IL-2 (PeproTech) at a density of 10^6^ cells per ml. Two days after stimulation, magnetic beads were removed, and T cells were ready for nucleofection. Edited T cells were cultured in X-vivo 15 medium supplemented with 5% human AB serum (Heat inactivated) and 200U/ml IL2. Medium was replaced every other day and T cells were maintained at a density of ∼10^6^ cells per ml. Unused isolated T cells were frozen in BamBanker freezing medium (Bulldog) and stored at −80(. If frozen T cells were used for activation and stimulation, T cell were rested in X-vivo 15 medium supplemented with 5% human AB serum for one day and then activated and stimulated as described above.

### Human T-cell nucleofection and DNA isolation

Nucleofection in T cells were performed using P3 Primary Cell 4D-Nucleofector^TM^ X KIT S (Lonza) according to the manufacturer’s instruction with minor modifications. Briefly, two days after initiating T cell activation and stimulation, ∼3-5 x 10^5^ T cells were collected and resuspended in 20 μl P3 buffer. 1 μg Cas9 mRNA and 1 μg gRNA for each gene alone or together with T4 DNA polymerase mRNA were added to the cells before nucleofection in a Lonza 4D-Nucleofector with pulse code EO-115. 80 μl X-vivo 15 medium with 5% human AB serum was added to the nucleofected cells before a 15 min recovery at 37 °C. Nucleofected T cells were plated at a density of ∼0.5–1 x 10^6^ cells/ml in X-vivo 15 medium supplemented with 5% human AB serum and 200U/mL IL-2 in 48-well plates. Edited T cells were replenished as needed to maintain a density of 10^6^ cells per ml. Four days after nucleofection, DNA were isolated from edited T cells using the DNeasy Blood and Tissue Kit (Qiagen) according to the manufacturer’s instruction.

### PCR amplicon preparation for deep sequencing

PCR amplicons of ∼300 bp were amplified using a GoTaq kit (Promega) with target-specific primers, separated on a 2% agarose gel, and purified with the MinElute Gel Extraction Kit (Qiagen). PCR amplicons were barcoded with the Nextera Flex Prep HT kit according to the manufacturer’s instruction and sequenced using the Miseq paired-end 150-cycle format by the Genome Technology Center Core Facility at New York University Langone Health. Primers for PCR are listed in **Supplemental table S2**.

### PCR amplicon preparation for PacBio sequencing

Multiplexed PacBio amplicon libraries preparation was performed using PacBio barcoded M13 primers and SMRTbell prep kit 3.0 (PacBio) according to the manufacturer’s instruction by the Genome Technology Center Core Facility at New York University Langone Health. Briefly, a two-step PCR process was required for generating barcoded amplicons for multiplexed PacBio sequencing. The first-round PCR was amplified using LA Taq DNA polymerase (Takara) with primers that contain a combination of universal and target-specific sequences. The first-round PCR products were cleaned with 2x AMPure PB beads (PacBio). Barcodes were incorporated into the first-round PCR product using universal sequences tailed with 17 bp M13 PacBio barcode sequences after the second round of PCR. Barcoded samples were then pooled as one sample for SMRTbell library construction. SMRTbell libraries were sequenced on the Sequel II and IIe systems. Primers for PacBio first-round PCR amplicons are listed in **Supplemental table S3**.

### Deep sequencing reads analysis

Unmapped paired-end deep sequencing reads were used as inputs into the CRISPResso2 tool to quantify the frequency of editing events(Pinello et al., 2016). The tool was run with default parameters (https://github.com/pinellolab/CRISPResso2).

### PacBio sequencing reads analysis

Raw PacBio data were demultiplexed with the corresponding barcode using the SMRTlink software to assign barcoded reads to each sample (smrtlink version: 8.0.0.80529, chemistry bundle: 8.0.0.778409, params: 8.0.0). Analysis of demultiplexed data was performed using PacBio tools distributed via Bioconda (https://github.com/PacificBiosciences/pbbioconda). Circular consensus sequences were converted to HiFi calls using the pbccs command and filtering for reads with support from at least three full-length subreads. The resulting fastq files were used as inputs to a custom python script that filtered for reads containing specific 20-bp index sequences at both the 5′ and 3′ regions of each read. Resulting filtered reads were mapped to the reference genome using minimap2 (-t 4 -ax splice --splice-flank=no -u no -G 5000). The genome coverage of the alignment files was calculated using the “bedtools genomecov -d -split - ibam” command. Normalized depth coverage was calculated by dividing the depth coverage at each position by the total number of reads (e.g., normalized depth coverage of G at position 10=100 x depth coverage at position 10/total number of reads). All downstream analyses were performed using custom Python scripts. The index sequences for each gene are listed in **Supplemental table S3**. The index sequences for each gene are listed in **Supplemental table S3**. **Droplet digital PCR**

Genomic DNA was isolated from T cells four days post-nucleofection using the DNeasy Blood and Tissue Kit (Qiagen) according to the manufacturer’s instruction. ddPCR reactions were assembled with ddPCR supermix for Probes (No dUTP, Bio-Rad), 900 nM of each primer, 250 nM of each probe, and 20 ng genome DNA. Droplets were formed using a Bio-Rad QX200 Droplet Generator following the manufacturer’s instruction and analyzed on a Bio-Rad QX200 Droplet reader. Data were analyzed with the QX manager Software (Bio-Rad), and thresholds were set manually based on the wells with untreated samples. The percentage of alleles with chromosome loss was calculated based on the ratio of droplets that had amplicon 2 (FAM+) to droplets that had amplicon 1 (VIC+). The equation utilized is as follows: % Chromosome loss = 100 x (1- [FAM]/[VIC]). The percentage of alleles with unintended edits was calculated based on the ratio of droplets that had amplicon 2 (FAM+) to droplets that had RPP30 (HEX+). The equation utilized is as follows: % Chromosome loss = 100 x (1- [FAM]/[HEX]). Primers and probes for ddPCR assay are listed in **Supplemental table S4**.

## Data availability

All data needed to evaluate the conclusions in the paper are present in the paper and/or the Supplementary information. Additional data related to this paper may be requested from the authors.

## Acknowledgements

This work was supported by grants from the Departmental Start-Up Grant, NYU Langone Health, and Kids Connect Charitable Fund.

