## Supplemental materials for "Preventing large deletions and chromosome loss in engineered human primary T cells by CasPlus with optimized guide RNAs"

by CasPlus with optimized guide RNA

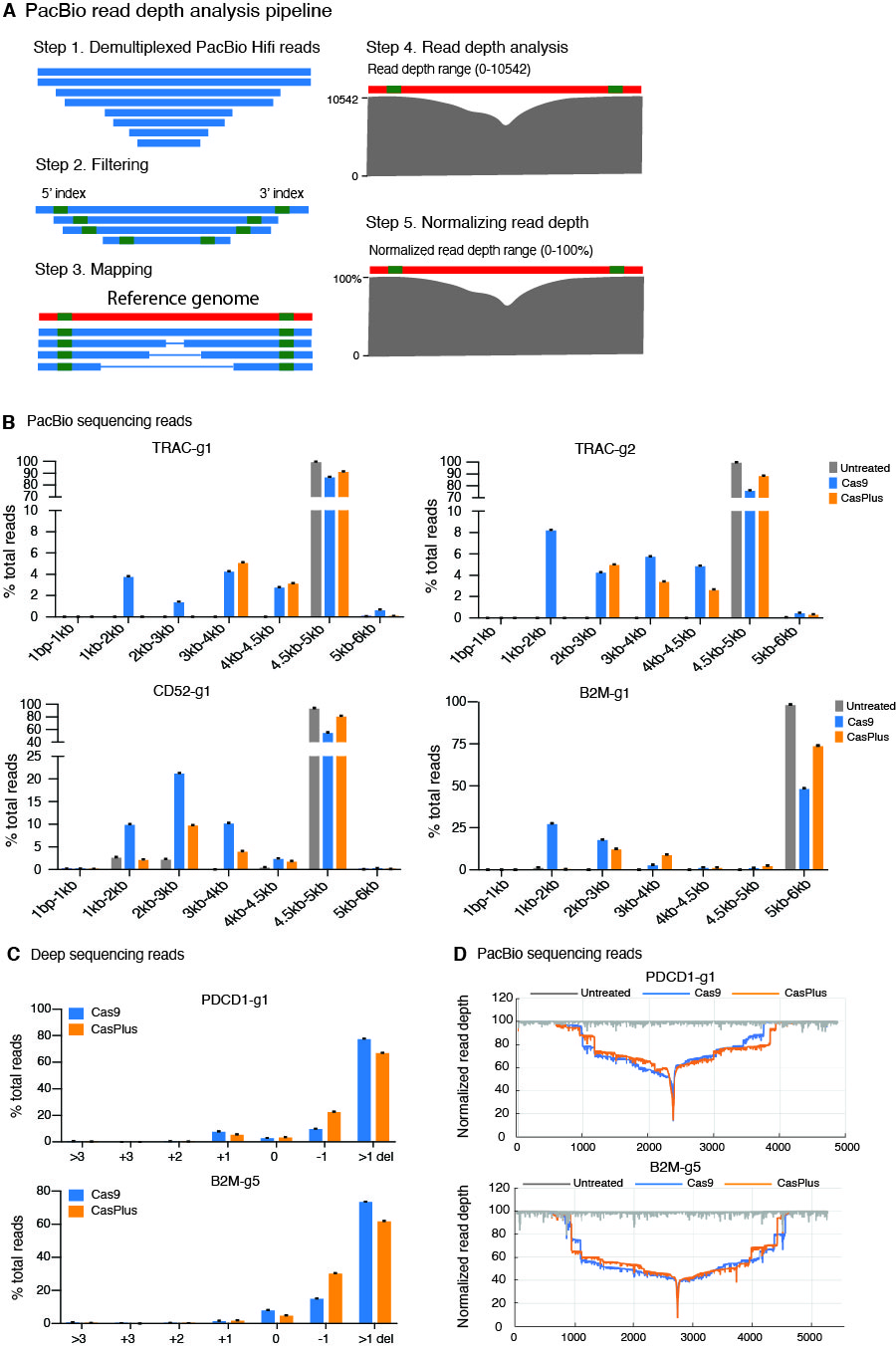

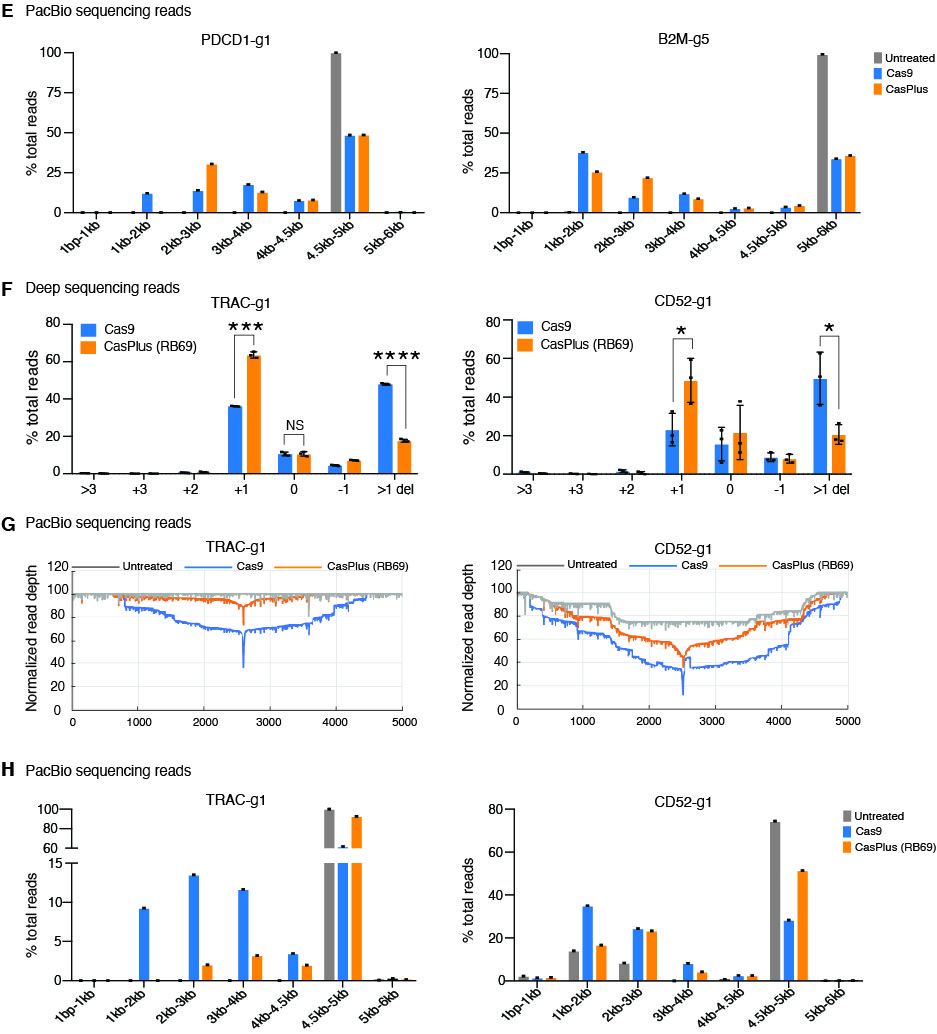
**Fig. S1. T4 and RB69 DNA polymerase mitigated large deletions in engineered human primary T cells, related to Figure 1.** (**A**) Schematics illustrating the pipeline for PacBio read depth analysis. Raw PacBio data were demultiplexed with the corresponding barcodes, and circular consensus sequences of each sample were converted to HiFi calls (Step 1). Demultiplexed PacBio HiFi reads were then used as inputs to a custom python script that filtered for reads containing specific 20-bp index sequences at both the 5′ and 3′ regions of each read (Step 2). The resulting filtered reads were mapped to the reference genome (Step 3) and genome coverage of the alignment files was calculated (Step 4). Normalized depth coverage was calculated by dividing the depth coverage at each position by the total number of reads (Step 5). (**B**) Quantification of PacBio reads with distinct sizes in samples described in **Fig. 1D**. (**C**) Frequencies of indels at PDCD1 and B2M loci in T cells nucleofected with PDCD1-g1 and B2M-g5, respectively, for Cas9 and CasPlus editing. Each dot represents one biological replicate. (**D**) Normalized PacBio reads depth at PDCD1 and B2M loci in untreated, Cas9-edited, and CasPlus-edited T cells with PDCD1-g1 and B2M-g5, respectively, for Cas9 and CasPlus editing. (**E**) Quantification of PacBio reads with distinct sizes in samples described in panel (**D**). (**F**) Frequencies of indels at TRAC and CD52 loci in T cells nucleofected with TRAC-g1 and CD52-g1, respectively, for Cas9 and RB69-mediated CasPlus editing. (**G**) Normalized PacBio reads depth at TRAC and CD52 loci in untreated, Cas9-edited, and CasPlus-edited T cells with TRAC-g1 and CD52-g1, respectively, for Cas9 and RB69-mediated CasPlus editing. (**H**) Quantification of PacBio reads with distinct sizes in samples described in panel (**G**).

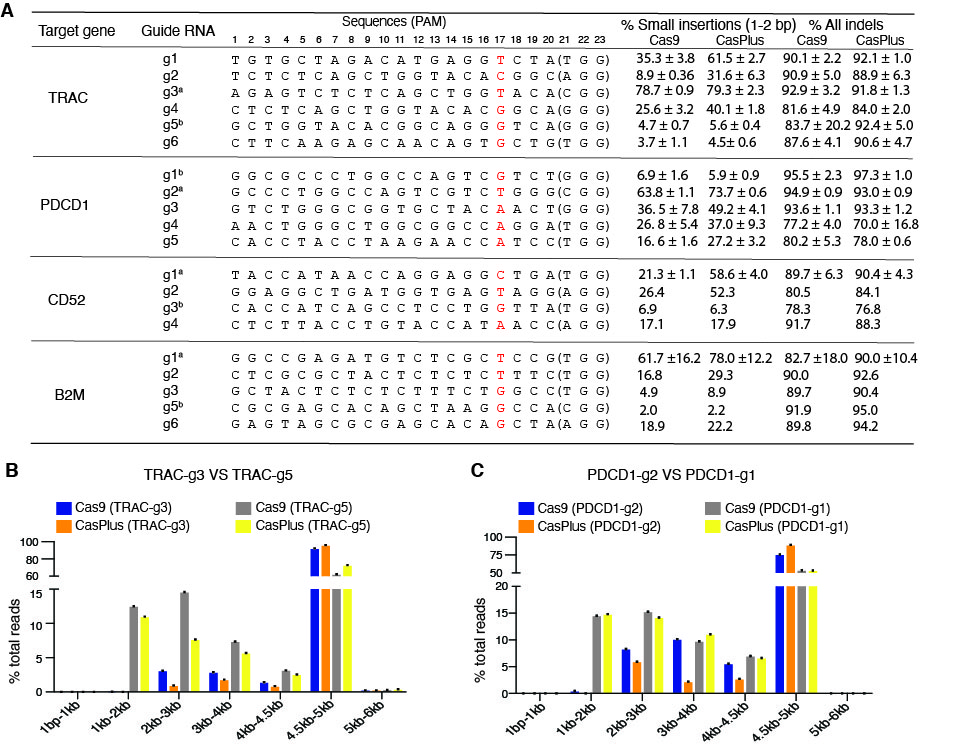

**Fig. S2. The frequencies of small insertions induced by Cas9 and CasPlus were gRNA context-dependent in engineered human primary T cells, related to Figure 2.** (**A**) Table demonstrating the sequences of the gRNAs designed to target genes *TRAC*, *PDCD1*, *CD52* and *B2M* and the quantification of total indels and small insertions induced by them in Cas9 and CasPlus editing within T cells. Nucleotides at position +17 were highlighted in red color. “a” labels the gRNA that was most efficient in inducing small insertions (1-2 bp) while “b” labels the gRNA that produced minimal small insertions. (**B**) Quantification of PacBio reads with distinct sizes in samples described in **Fig. 2B**. (**C**) Quantification of PacBio reads with distinct sizes in samples descripted in **Fig. 2E**.

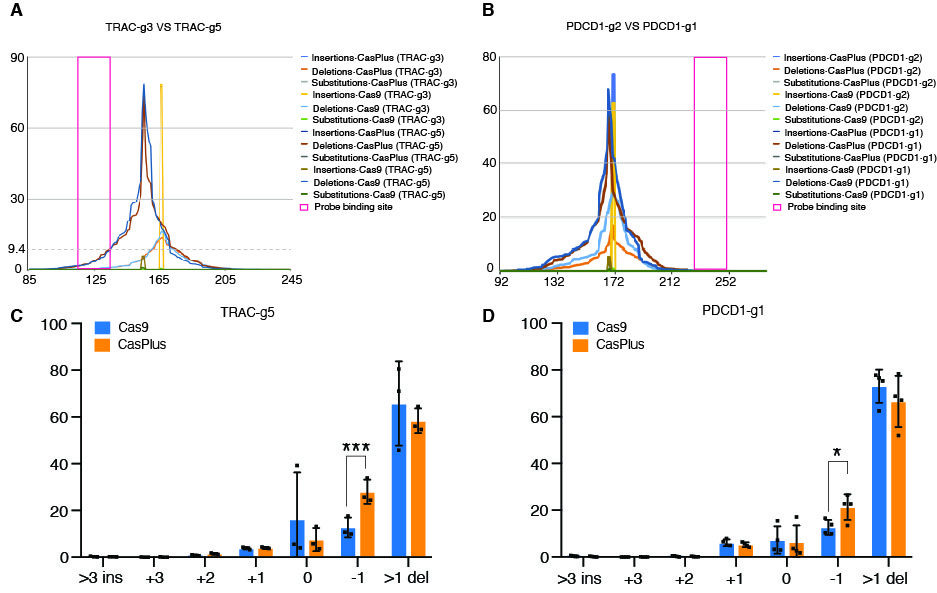

**Fig. S3. The activity of CasPlus editing in mitigating chromosome loss is associated with its enhancement in small insertions, but not deletions, related to Figure 3. (A**) Distribution of Cas9- and CasPlus-induced indels (insertions, deletions, and substitutions) at the TRAC locus from deep sequencing of T cells with TRAC-g3 or TRAC-g5 for Cas9 and CasPlus editing. **(B**) Distribution of Cas9- and CasPlus-induced indels (insertions, deletions, and substitutions) at the PDCD1 locus from deep sequencing of T cells with PDCD1-g2 or PDCD1-g1 for Cas9 and CasPlus editing. (**C-D**) Frequencies of indels induced by TRAC-g5 (**C**) or PDCD1-g1 (**D**) for Cas9 and CasPlus editing within T cells.

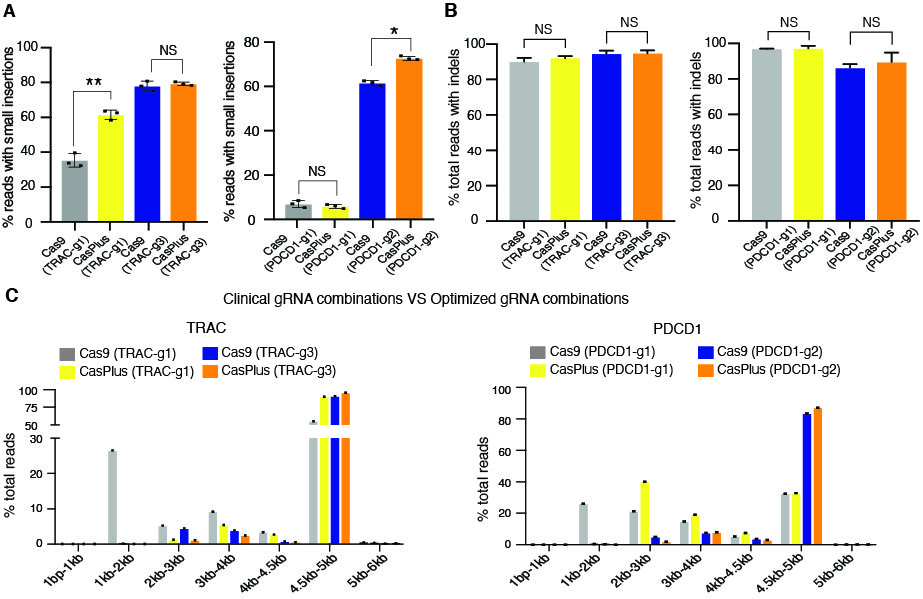

**Fig. S4. Optimized gRNA combinations resulted in higher small insertions compared to clinically used gRNA combination in T cells**. (**A-B**). Quantification of total small insertions (**A**) and total indels (**B**) at the *TRAC* and *PDCD1* individual sites in T cells with TRAC-g1 and PDCD1-g1 or TRAC-g3 and PDCD1-g2 for Cas9 and CasPlus editing. (**C**) Quantification of PacBio reads with distinct sizes in samples described in **Fig. 4B.**

**Supplemental table S1.** Quantification of PacBio reads with distinct sizes.

| Size | TRAC-g1 (Reads #) | | | TRAC-g1 (% Reads) | | |
| --- | --- | --- | --- | --- | --- | --- |
|  | Untreated | Cas9 | CasPlus | Untreated | Cas9 | CasPlus |
| 1bp-1kb | 0 | 0 | 0 | 0 | 0 | 0 |
| 1kb-2kb | 0 | 501 | 0 | 0 | 3.812495 | 0 |
| 2kb-3kb | 0 | 187 | 2 | 0 | 1.423027 | 0.009615 |
| 3kb-4kb | 1 | 565 | 1068 | 0.00249 | 4.299521 | 5.134615 |
| 4kb-4.5kb | 1 | 368 | 659 | 0.00249 | 2.800396 | 3.168269 |
| 4.5kb-5kb | 40122 | 11430 | 19054 | 99.9228 | 86.97968 | 91.60577 |
| 5kb-6kb | 29 | 90 | 17 | 0.072224 | 0.684879 | 0.081731 |
| Total | 40153 | 13141 | 20800 | 100 | 100 | 100 |
| Size | TRAC-g2 (Reads #) | | | TRAC-g2 (% Reads) | | |
|  | Untreated | Cas9 | CasPlus | Untreated | Cas9 | CasPlus |
| 1bp-1kb | 0 | 0 | 0 | 0 | 0 | 0 |
| 1kb-2kb | 0 | 2248 | 0 | 0 | 8.254388 | 0 |
| 2kb-3kb | 0 | 1169 | 1286 | 0 | 4.292429 | 5.02599 |
| 3kb-4kb | 1 | 1573 | 876 | 0.00249 | 5.775868 | 3.423614 |
| 4kb-4.5kb | 1 | 1334 | 683 | 0.00249 | 4.898289 | 2.669324 |
| 4.5kb-5kb | 40122 | 20775 | 22653 | 99.9228 | 76.28332 | 88.53324 |
| 5kb-6kb | 29 | 135 | 89 | 0.072224 | 0.495704 | 0.347833 |
| Total | 40153 | 27234 | 25587 | 100 | 100 | 100 |
| Size | CD52-g1 (Reads #) | | | CD52-g1 (% Reads) | | |
|  | Untreated | Cas9 | CasPlus | Untreated | Cas9 | CasPlus |
| 1bp-1kb | 115 | 66 | 52 | 0.193199 | 0.192532 | 0.136853 |
| 1kb-2kb | 1648 | 3441 | 849 | 2.768631 | 10.03792 | 2.234387 |
| 2kb-3kb | 1386 | 7307 | 3733 | 2.328473 | 21.31564 | 9.82446 |
| 3kb-4kb | 10 | 3523 | 1546 | 0.0168 | 10.27713 | 4.068742 |
| 4kb-4.5kb | 305 | 845 | 716 | 0.512398 | 2.464994 | 1.884359 |
| 4.5kb-5kb | 56033 | 18997 | 31059 | 94.13514 | 55.41715 | 81.74066 |
| 5kb-6kb | 27 | 101 | 42 | 0.04536 | 0.294632 | 0.110535 |
| Total | 59524 | 34280 | 37997 | 100 | 100 | 100 |
| Size | B2M-g1 (Reads #) | | | B2M-g1 (% Reads) | | |
|  | Untreated | Cas9 | CasPlus | Untreated | Cas9 | CasPlus |
| 1bp-1kb | 0 | 0 | 0 | 0 | 0 | 0 |
| 1kb-2kb | 63 | 4947 | 32 | 0.40478 | 27.67243 | 0.319521 |
| 2kb-3kb | 4 | 3235 | 1247 | 0.0257 | 18.09588 | 12.45132 |
| 3kb-4kb | 1 | 533 | 916 | 0.006425 | 2.981485 | 9.146281 |
| 4kb-4.5kb | 0 | 268 | 148 | 0 | 1.499133 | 1.477783 |
| 4.5kb-5kb | 1 | 206 | 258 | 0.006425 | 1.152319 | 2.576136 |
| 5kb-6kb | 15495 | 8688 | 7414 | 99.55667 | 48.59876 | 74.02896 |
| Total | 15564 | 17877 | 10015 | 100 | 100 | 100 |
| Size | PDCD1-g1 (Reads #) | | | PDCD1-g1 (% Reads) | | |
|  | Untreated | Cas9 | CasPlus | Untreated | Cas9 | CasPlus |
| 1bp-1kb | 0 | 0 | 0 | 0 | 0 | 0 |
| 1kb-2kb | 2 | 1019 | 8 | 0.006355 | 12.19629 | 0.13369 |
| 2kb-3kb | 1 | 1163 | 1823 | 0.003177 | 13.91981 | 30.46457 |
| 3kb-4kb | 1 | 1471 | 772 | 0.003177 | 17.60622 | 12.90107 |
| 4kb-4.5kb | 1 | 634 | 474 | 0.003177 | 7.588271 | 7.921123 |
| 4.5kb-5kb | 31460 | 4050 | 2906 | 99.96187 | 48.47397 | 48.56283 |
| 5kb-6kb | 7 | 18 | 1 | 0.022242 | 0.21544 | 0.016711 |
| Total | 31472 | 8355 | 5984 | 100 | 100 | 100 |
| Size | B2M-g5 (Reads #) | | | B2M-g5 (% Reads) | | |
|  | Untreated | Cas9 | CasPlus | Untreated | Cas9 | CasPlus |
| 1bp-1kb | 0 | 0 | 0 | 0 | 0 | 0 |
| 1kb-2kb | 63 | 4433 | 2727 | 0.40478 | 37.98629 | 25.69732 |
| 2kb-3kb | 4 | 1140 | 2334 | 0.0257 | 9.768638 | 21.99397 |
| 3kb-4kb | 1 | 1400 | 936 | 0.006425 | 11.99657 | 8.820204 |
| 4kb-4.5kb | 0 | 320 | 316 | 0 | 2.742074 | 2.977761 |
| 4.5kb-5kb | 1 | 411 | 484 | 0.006425 | 3.521851 | 4.560874 |
| 5kb-6kb | 15495 | 3966 | 3815 | 99.55667 | 33.98458 | 35.94987 |
| Total | 15564 | 11670 | 10612 | 100 | 99.9999999 | 100 |
| Size | TRAC-g3 (Reads #)  (Single gene-edited T cells) | | | TRAC-g3 (% Reads)  (Single gene-edited T cells) | | |
|  | Untreated | Cas9 | CasPlus | Untreated | Cas9 | CasPlus |
| 1bp-1kb | 0 | 0 | 0 | 0 | 0 | 0 |
| 1kb-2kb | 0 | 2 | 0 | 0 | 0.023521 | 0 |
| 2kb-3kb | 0 | 263 | 91 | 0 | 3.093026 | 0.947423 |
| 3kb-4kb | 1 | 245 | 173 | 0.00249 | 2.881336 | 1.801145 |
| 4kb-4.5kb | 1 | 121 | 81 | 0.00249 | 1.423027 | 0.843311 |
| 4.5kb-5kb | 40122 | 7854 | 9240 | 99.9228 | 92.3674 | 96.1999 |
| 5kb-6kb | 29 | 18 | 20 | 0.072224 | 0.21169 | 0.208225 |
| Total | 40153 | 8503 | 9605 | 100 | 100 | 100 |
| Size | TRAC-g5 (Reads #)  (Single gene-edited T cells) | | | TRAC-g5 (% Reads)  (Single gene-edited T cells) | | |
|  | Untreated | Cas9 | CasPlus | Untreated | Cas9 | CasPlus |
| 1bp-1kb | 0 | 0 | 0 | 0 | 0 | 0 |
| 1kb-2kb | 0 | 799 | 1025 | 0 | 12.51174 | 10.99432 |
| 2kb-3kb | 0 | 932 | 713 | 0 | 14.59443 | 7.647753 |
| 3kb-4kb | 1 | 471 | 531 | 0.00249 | 7.375509 | 5.695592 |
| 4kb-4.5kb | 1 | 201 | 237 | 0.00249 | 3.14751 | 2.5421 |
| 4.5kb-5kb | 40122 | 3965 | 6781 | 99.9228 | 62.08894 | 72.7341 |
| 5kb-6kb | 29 | 18 | 36 | 0.072224 | 0.281867 | 0.386142 |
| Total | 40153 | 6386 | 9323 | 100 | 100 | 100 |
| Size | PDCD1-g1 (Reads #)  (Single gene-edited T cells) | | | PDCD1-g1 (% Reads)  (Single gene-edited T cells) | | |
|  | Untreated | Cas9 | CasPlus | Untreated | Cas9 | CasPlus |
| 1bp-1kb | 0 | 0 | 0 | 0 | 0 | 0 |
| 1kb-2kb | 0 | 58 | 47 | 0 | 14.5 | 14.8265 |
| 2kb-3kb | 0 | 61 | 45 | 0 | 15.25 | 14.19558 |
| 3kb-4kb | 0 | 39 | 35 | 0 | 9.75 | 11.04101 |
| 4kb-4.5kb | 0 | 28 | 21 | 0 | 7 | 6.624606 |
| 4.5kb-5kb | 652 | 214 | 169 | 99.6941896 | 53.5 | 53.3123 |
| 5kb-6kb | 2 | 0 | 0 | 0.30581 | 0 | 0 |
| Total | 654 | 400 | 317 | 100 | 100 | 100 |
| Size | PDCD1-g2 (Reads #)  (Single gene-edited T cells) | | | PDCD1-g2 (% Reads)  (Single gene-edited T cells) | | |
|  | Untreated | Cas9 | CasPlus | Untreated | Cas9 | CasPlus |
| 1bp-1kb | 0 | 0 | 0 | 0 | 0 | 0 |
| 1kb-2kb | 0 | 1 | 0 | 0 | 0.460829 | 0 |
| 2kb-3kb | 0 | 18 | 24 | 0 | 8.294931 | 5.91133 |
| 3kb-4kb | 0 | 22 | 9 | 0 | 10.13825 | 2.216749 |
| 4kb-4.5kb | 0 | 12 | 11 | 0 | 5.529954 | 2.70936 |
| 4.5kb-5kb | 652 | 164 | 362 | 99.6941896 | 75.57604 | 89.16256 |
| 5kb-6kb | 2 | 0 | 0 | 0.30581 | 0 | 0 |
| Total | 654 | 217 | 406 | 100 | 100 | 100 |
| Size | TRAC-g1 (Reads #)  (Multiple gene-edited T cells) | | | TRAC-g1 (% Reads)  (Multiple gene-edited T cells) | | |
|  | Untreated | Cas9 | CasPlus | Untreated | Cas9 | CasPlus |
| 1bp-1kb | 0 | 0 | 0 | 0 | 0 | 0 |
| 1kb-2kb | 0 | 8483 | 24 | 0 | 26.57165 | 0.092226 |
| 2kb-3kb | 0 | 1681 | 319 | 0 | 5.265466 | 1.225839 |
| 3kb-4kb | 1 | 2942 | 1412 | 0.00249 | 9.215348 | 5.425969 |
| 4kb-4.5kb | 1 | 1071 | 697 | 0.00249 | 3.354738 | 2.6784 |
| 4.5kb-5kb | 40122 | 17591 | 23481 | 99.9228 | 55.10102 | 90.23172 |
| 5kb-6kb | 29 | 157 | 90 | 0.072224 | 0.491778 | 0.345848 |
| Total | 40153 | 31925 | 26023 | 100 | 100 | 100 |
| Size | TRAC-g3 (Reads #)  (Multiple gene-edited T cells) | | | TRAC-g3 (% Reads)  (Multiple gene-edited T cells) | | |
|  | Untreated | Cas9 | CasPlus | Untreated | Cas9 | CasPlus |
| 1bp-1kb | 0 | 0 | 0 | 0 | 0 | 0 |
| 1kb-2kb | 0 | 2 | 4 | 0 | 0.005079 | 0.011944 |
| 2kb-3kb | 0 | 1756 | 358 | 0 | 4.459796 | 1.068944 |
| 3kb-4kb | 1 | 1528 | 818 | 0.00249 | 3.880733 | 2.442447 |
| 4kb-4.5kb | 1 | 301 | 172 | 0.00249 | 0.764464 | 0.513571 |
| 4.5kb-5kb | 40122 | 35747 | 32063 | 99.9228 | 90.78834 | 95.73617 |
| 5kb-6kb | 29 | 40 | 76 | 0.072224 | 0.10159 | 0.226927 |
| Total | 40153 | 39374 | 33491 | 100 | 100 | 100 |
| Size | PDCD1-g1 (Reads #)  (Multiple gene-edited T cells) | | | PDCD1-g1 (% Reads)  (Multiple gene-edited T cells) | | |
|  | Untreated | Cas9 | CasPlus | Untreated | Cas9 | CasPlus |
| 1bp-1kb | 0 | 1 | 0 | 0 | 0.013452 | 0 |
| 1kb-2kb | 2 | 1938 | 48 | 0.006355 | 26.06941 | 0.516018 |
| 2kb-3kb | 1 | 1591 | 3729 | 0.003177 | 21.40167 | 40.08815 |
| 3kb-4kb | 1 | 1099 | 1768 | 0.003177 | 14.78343 | 19.00667 |
| 4kb-4.5kb | 1 | 390 | 697 | 0.003177 | 5.246166 | 7.493012 |
| 4.5kb-5kb | 31460 | 2413 | 3051 | 99.96187 | 32.45897 | 32.7994 |
| 5kb-6kb | 7 | 2 | 9 | 0.022242 | 0.026903 | 0.096753 |
| Total | 31472 | 7434 | 9302 | 100 | 100 | 100 |
| Size | PDCD1-g2 (Reads #)  (Multiple gene-edited T cells) | | | PDCD1-g2 (% Reads)  (Multiple gene-edited T cells) | | |
|  | Untreated | Cas9 | CasPlus | Untreated | Cas9 | CasPlus |
| 1bp-1kb | 0 | 0 | 0 | 0 | 0 | 0 |
| 1kb-2kb | 2 | 46 | 3 | 0.006355 | 0.381774 | 0.017865 |
| 2kb-3kb | 1 | 598 | 318 | 0.003177 | 4.963067 | 1.893646 |
| 3kb-4kb | 1 | 900 | 1324 | 0.003177 | 7.4695 | 7.884237 |
| 4kb-4.5kb | 1 | 436 | 512 | 0.003177 | 3.618558 | 3.048889 |
| 4.5kb-5kb | 31460 | 10053 | 14617 | 99.96187 | 83.43431 | 87.04222 |
| 5kb-6kb | 7 | 16 | 19 | 0.022242 | 0.132791 | 0.113142 |
| Total | 31472 | 12049 | 16793 | 100 | 100 | 100 |

**Supplemental table S2**. gRNA and primer sequences for deep sequencing.

| gRNA_name | gRNA_sequence | gRNA_PAM | DS_primer_f | DS_primer_r |
| --- | --- | --- | --- | --- |
| TRAC-g1 | TGTGCTAGACATGAGGTCTA | TGG | agatatccagaaccctgacc | cctgaagcaaggaaacag |
| TRAC-g2 | TCTCTCAGCTGGTACACGGC | AGG | cgtataaagcatgagaccgtg | cagcactgttgctcttgaag |
| TRAC-g3 | AGAGTCTCTCAGCTGGTACA | CGG | cgtataaagcatgagaccgtg | cagcactgttgctcttgaag |
| TRAC-g4 | CTCTCAGCTGGTACACGGCA | GGG | cgtataaagcatgagaccgtg | cagcactgttgctcttgaag |
| TRAC-g5 | GCTGGTACACGGCAGGGTCA | GGG | cgtataaagcatgagaccgtg | cagcactgttgctcttgaag |
| TRAC-g6 | CTTCAAGAGCAACAGTGCTG | TGG | agatatccagaaccctgacc | cctgaagcaaggaaacag |
| PDCD1-g1 | GGCGCCCTGGCCAGTCGTCT | GGG | ctgccagggccaggcccg | acgtggatgtggaggaagag |
| PDCD1-g2 | CCCTGGCCAGTCGTCTGGG | CGG | ctgccagggccaggcccg | acgtggatgtggaggaagag |
| PDCD1-g3 | TCTGGGCGGTGCTACAACT | GGG | ctgccagggccaggcccg | acgtggatgtggaggaagag |
| PDCD1-g4 | AACTGGGCTGGCGGCCAGGA | TGG | ctgccagggccaggcccg | acgtggatgtggaggaagag |
| PDCD1-g5 | CACCTACCTAAGAACCATCC | TGG | ctgccagggccaggcccg | acgtggatgtggaggaagag |
| CD52-g1 | TACCATAACCAGGAGGCTGA | TGG | ttcctctccctacctcacc | gttttcctgttggagtcc |
| CD52-g2 | GGAGGCTGATGGTGAGTAGG | AGG | ttcctctccctacctcacc | gttttcctgttggagtcc |
| CD52-g3 | CACCATCAGCCTCCTGGTTA | TGG | ttcctctccctacctcacc | gttttcctgttggagtcc |
| CD52-g4 | CTCTTACCTGTACCATAACC | AGG | ttcctctccctacctcacc | gttttcctgttggagtcc |
| B2M-g1 | GGCCGAGATGTCTCGCTCCG | TGG | taacctggcactgcgtcgct | ccaccaaggagaacttggag |
| B2M-g2 | CTCGCGCTACTCTCTCTTTC | TGG | taacctggcactgcgtcgct | ccaccaaggagaacttggag |
| B2M-g3 | GCTACTCTCTCTTTCTGGCC | TGG | taacctggcactgcgtcgct | ccaccaaggagaacttggag |
| B2M-g5 | CGCGAGCACAGCTAAGGCCA | CGG | taacctggcactgcgtcgct | ccaccaaggagaacttggag |
| B2M-g6 | GAGTAGCGCGAGCACAGCTA | AGG | taacctggcactgcgtcgct | ccaccaaggagaacttggag |

**Supplemental table S3**. Primer and index sequences for PacBio sequencing and analysis.

|  | PacBio_1st_round_f | PacBio_1st_round_r | 5' index | 3' index |
| --- | --- | --- | --- | --- |
| TRAC-g1 | /5AmMC6/GTAAAACGACGGCCAGTTTGGCAAAGGAACCAGAGTTTCCAC | /5AmMC6/CAGGAAACAGCTATGACAAGGCAAAACCTCTTGGCTGTCC | aaaaaaattaagtacctcta | ctgcccaagaactaggaggt |
| TRAC-g2 | /5AmMC6/GTAAAACGACGGCCAGTTTGGCAAAGGAACCAGAGTTTCCAC | /5AmMC6/CAGGAAACAGCTATGACAAGGCAAAACCTCTTGGCTGTCC | aaaaaaattaagtacctcta | ctgcccaagaactaggaggt |
| CD52-g1 | /5AmMC6/GTAAAACGACGGCCAGTAAAGAGGGTGATGGATGGCTGGGG | /5AmMC6/CAGGAAACAGCTATGACTCCCCACTGCAAGTACAAGGGTAAC | actcctccctcttctctccc | cccagctgggggtcaataaa |
| B2M-g1 | /5AmMC6/GTAAAACGACGGCCAGgggtggagtggctggatattatgg | /5AmMC6/CAGGAAACAGCTATGACtagaccatccatggggaagtggg | aaaaaaaggccgctattaat | aaattcaaacccagcctgtc |
| PDCD1-g1 | /5AmMC6/GTAAAACGACGGCCAGgggacaccgtatgtgtttgg | /5AmMC6/CAGGAAACAGCTATGACATCCTGCACGTCCAGCCCTCAGTCG | ctccctgtgcccgcccccta | ttagtgacagggtcttgctc |
| B2M-g5 | /5AmMC6/GTAAAACGACGGCCAGgggtggagtggctggatattatgg | /5AmMC6/CAGGAAACAGCTATGACtagaccatccatggggaagtggg | aaaaaaaggccgctattaat | aaattcaaacccagcctgtc |
| TRAC-g3 | /5AmMC6/GTAAAACGACGGCCAGTTTGGCAAAGGAACCAGAGTTTCCAC | /5AmMC6/CAGGAAACAGCTATGACAAGGCAAAACCTCTTGGCTGTCC | aaaaaaattaagtacctcta | ctgcccaagaactaggaggt |
| TRAC-g5 | /5AmMC6/GTAAAACGACGGCCAGTTTGGCAAAGGAACCAGAGTTTCCAC | /5AmMC6/CAGGAAACAGCTATGACAAGGCAAAACCTCTTGGCTGTCC | aaaaaaattaagtacctcta | ctgcccaagaactaggaggt |
| PDCD1-g2 | /5AmMC6/GTAAAACGACGGCCAGgggacaccgtatgtgtttgg | /5AmMC6/CAGGAAACAGCTATGACATCCTGCACGTCCAGCCCTCAGTCG | ctccctgtgcccgcccccta | ttagtgacagggtcttgctc |

**Supplemental table S4**. Primer and probe sequences for ddPCR.

| gRNA | ddPCR_  a1_primer_f | ddPCR_  a1_primer_r | ddPCR_  a1_probe | ddPCR_  a2_primer_f | ddPCR_  a2_primer_r | ddPCR_  a2_probe |
| --- | --- | --- | --- | --- | --- | --- |
| TRAC-g1 | agcctcagtctctccaactg | tgactgcgtgagactgactt | tcctgcctgcctgcctttgc | tgattctcaaacaaatgtgtcac | aaagctgcccttacctg | cgccttcaacaacagcattattcca |
| TRAC-g3 | agcctcagtctctccaactg | tgactgcgtgagactgactt | tcctgcctgcctgcctttgc | gggcaaagagggaaatgaga | tgtgacacatttgtttgagaatc | tgggacaagaggatcagggt |
| TRAC-g5 | agcctcagtctctccaactg | tgactgcgtgagactgactt | tcctgcctgcctgcctttgc | gggcaaagagggaaatgaga | tgtgacacatttgtttgagaatc | tgggacaagaggatcagggt |
| PDCD1-g1 | agtgaggaccaaggatgc | tcctgggttcctctctg | cctgcctgcccaggagcaa | gactgagggtggaaggtc | cctgagcagtggagaag | cctggctctgggacacctgacc |
| PDCD1-g2 | agtgaggaccaaggatgc | tcctgggttcctctctg | cctgcctgcccaggagcaa | gactgagggtggaaggtc | cctgagcagtggagaag | cctggctctgggacacctgacc |
